# Lessons from simple marine models on the bacterial regulation of eukaryotic development

**DOI:** 10.1101/211797

**Authors:** Arielle Woznica, Nicole King

## Abstract

Molecular cues from environmental bacteria influence important developmental decisions in diverse marine eukaryotes. Yet, relatively little is understood about the mechanisms underlying these interactions, in part because marine ecosystems are dynamic and complex. With the help of simple model systems, including the choanoflagellate *Salpingoeca rosetta*, we have begun to uncover the bacterial cues that shape eukaryotic development in the ocean. Here, we review how diverse bacterial cues – from lipids to macromolecules – regulate development in marine eukaryotes. It is becoming clear that there are networks of chemical information circulating in the ocean, with both eukaryotes and bacteria acting as nodes; one eukaryote can precisely respond to cues from several diverse environmental bacteria, and a single environmental bacterium can regulate the development of different eukaryotes.

**Highlights:**

- Cues from environmental bacteria influence the development of many marine eukaryotes
- The molecular cues produced by environmental bacteria are structurally diverse
- Eukaryotes can respond to many different environmental bacteria
- Some environmental bacteria act as “information hubs” for diverse eukaryotes
- Experimentally tractable systems, like the choanoflagellate *S. rosetta*, promise to reveal molecular mechanisms underlying these interactions

## Introduction

Eukaryotes evolved over two billion years ago in a world dominated by prokaryotes and have lived in close association with bacteria ever since. It has become increasingly clear that bacteria not only act as competitors and pathogens, but also promote proper health and development in eukaryotes [1,2]. Growing attention has focused on how stably associated bacteria (e.g. the microbiome) shape many aspects of eukaryotic development, from root nodule development in legumes [3] to light organ morphogenesis in the Hawaiin bobtail squid [4], and even immune system development in vertebrates [5]. Yet, bacteria in the microbiome are not the only bacteria influencing eukaryotic development. Although often overlooked, free-living environmental bacteria also provide cues that regulate essential developmental processes in diverse eukaryotes.

Many examples of interactions between environmental bacteria and eukaryotes stem from marine ecosystems, where bacterial cues elicit developmental transitions in organisms as diverse as algae and animals. Nonetheless, few of these bacterial-eukaryotic interactions are understood in molecular detail, in part because marine environments are host to dynamic and diverse bacterial communities. While it is challenging to decipher specific interactions in such complex ecosystems, the lessons learned are likely to extend to interactions between eukaryotes and other bacterial communities, such as those in the gut and soil.

Simple model systems are beginning to reveal how environmental bacteria shape eukaryotic development in the ocean. Important features of these models that facilitate the identification of molecules underlying bacterial-eukaryotic interactions include: (1) the ability to grow and manipulate both the bacteria and the eukaryote in the lab, and (2) a clear and quantifiable response of the eukaryote to a single bacterium. Here we review mechanisms by which environmental bacteria regulate the development of choanoflagellates and other marine eukaryotes to illustrate how, and explore why, important eukaryotic developmental decisions rely on cues from specific environmental bacteria.

## A choanoflagellate model for bacterial-eukaryotic interactions

One of the closest living relatives of animals, the choanoflagellate *Salpingoeca rosetta*, has emerged as an attractive model for investigating how environmental bacteria shape eukaryotic cell biology and life history. Choanoflagellates are unicellular and colony-forming microeukaryotes that live in diverse aquatic environments [6]. Every choanoflagellate cell bears an apical “collar complex” – a single flagellum surrounded by a feeding collar composed of actin-filled microvilli – that it uses to capture and phagocytose bacterial prey. Importantly, the collar complex and its role in mediating interactions with bacteria are conserved among choanoflagellates and animals [6–8].

However, choanoflagellates do not just eat bacteria, but they also undergo key life history transitions in response to molecular cues secreted by environmental bacteria.

### A network of bacterial lipids flips a developmental switch in S. rosetta

In many choanoflagellates, including the emerging model choanoflagellate *S. rosetta*, a solitary cell can develop into a multicellular “rosette” colony through serial rounds of oriented cell division, with the sister cells remaining stably adherent [9,10] (Figure 1a). Although *S. rosetta* was isolated from the ocean as a rosette, early laboratory cultures proliferated primarily in the unicellular form, producing rosettes infrequently and unpredictably. A set of unexpected observations revealed that *Algoriphagus machipongonensis,* an environmental bacterium that had been co-isolated with the choanoflagellate and persisted in laboratory cultures at very low densities, could induce robust and uniform rosette development in *S. rosetta* when grown at higher densities [11].

**Figure 1.**
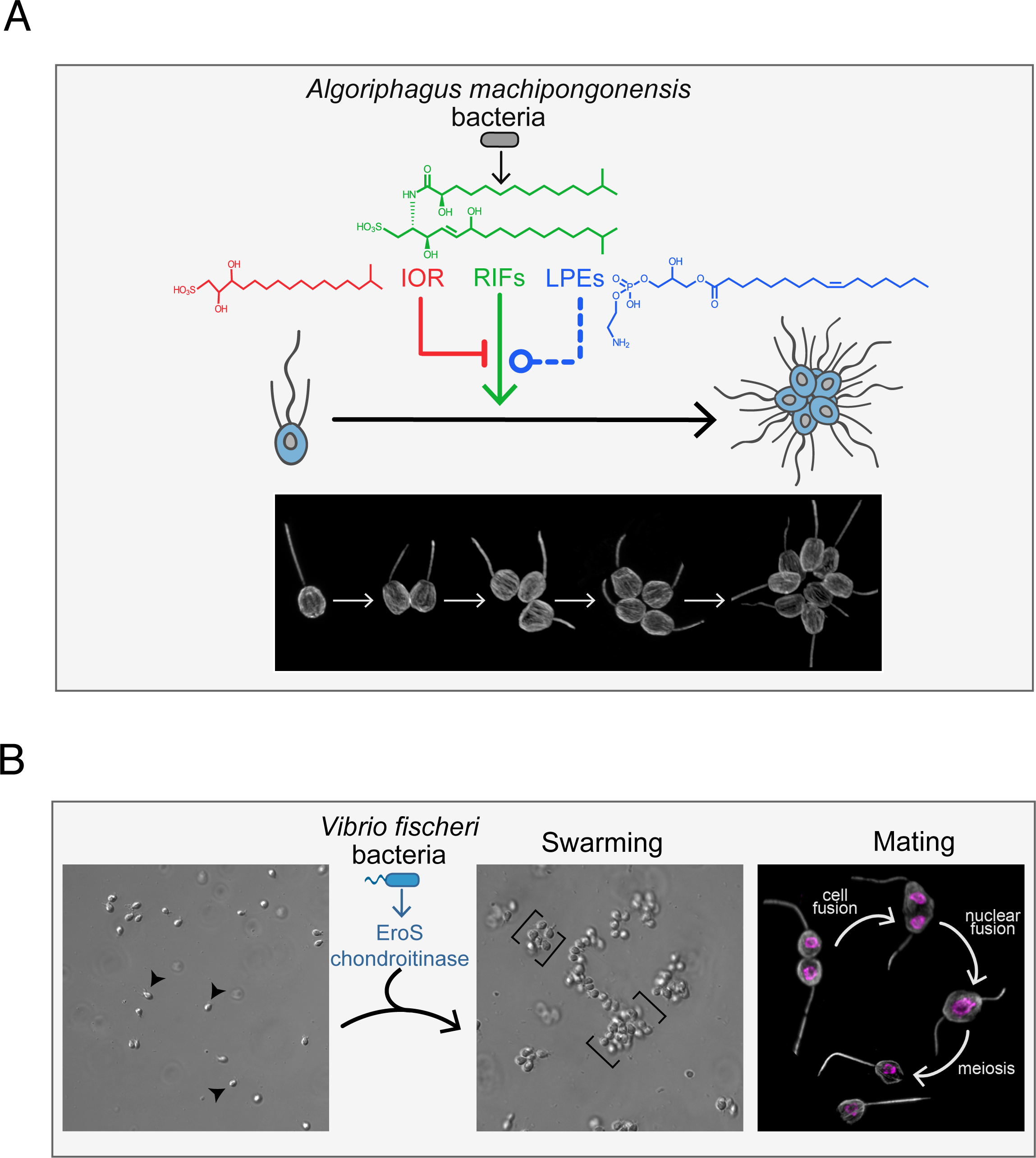
Bacteria regulate rosette development and sexual reproduction in the choanoflagellate, *S. rosetta*. **(A)** *Algoriphagus machipongonensis* bacteria regulate the development of *S. rosetta* from a solitary cell into a multicellular “rosette” colony through serial rounds of cell division*. Algoriphagus* produces three classes of lipids – sulfonolipids (RIFs), lysophosphatidylethanolamines (LPEs), and a capnine (IOR-1)– that interact to alternately induce, enhance, or inhibit rosette development. While the sulfonolipid RIFs are sufficient to initiate rosette development in *S. rosetta*, they require the synergistic enhancing activity of the LPEs for robust rosette development. *Algoriphagus* also produces the inhibitory IOR-1 that inhibits the RIFs, but cannot overcome the synergistic inducing activity of the RIFs + LPEs. Immunofluorescence images illustrate stages of *S. rosetta* rosette development; tubulin staining (grey) highlights the cell body and apical flagellum. **(B)** *Vibrio fischeri* bacteria induce sexual reproduction in *S. rosetta*. EroS, a chondroitin lyase secreted by *V. fischeri*, triggers solitary *S. rosetta* cells (arrows) to form large swarms (brackets) through cell aggregation. During swarming, *S. rosetta* cells pair off and mate, a process that involves the cell and nuclear fusion of two haploid cells into one diploid cell, followed by meiosis to generate haploid progeny. Immunofluorescence images depict mating stages in *S. rosetta*; tubulin staining (grey) highlights the cell body and apical flagellum, and Hoechst staining (magenta) highlights the nucleus.

**Figure 2.**
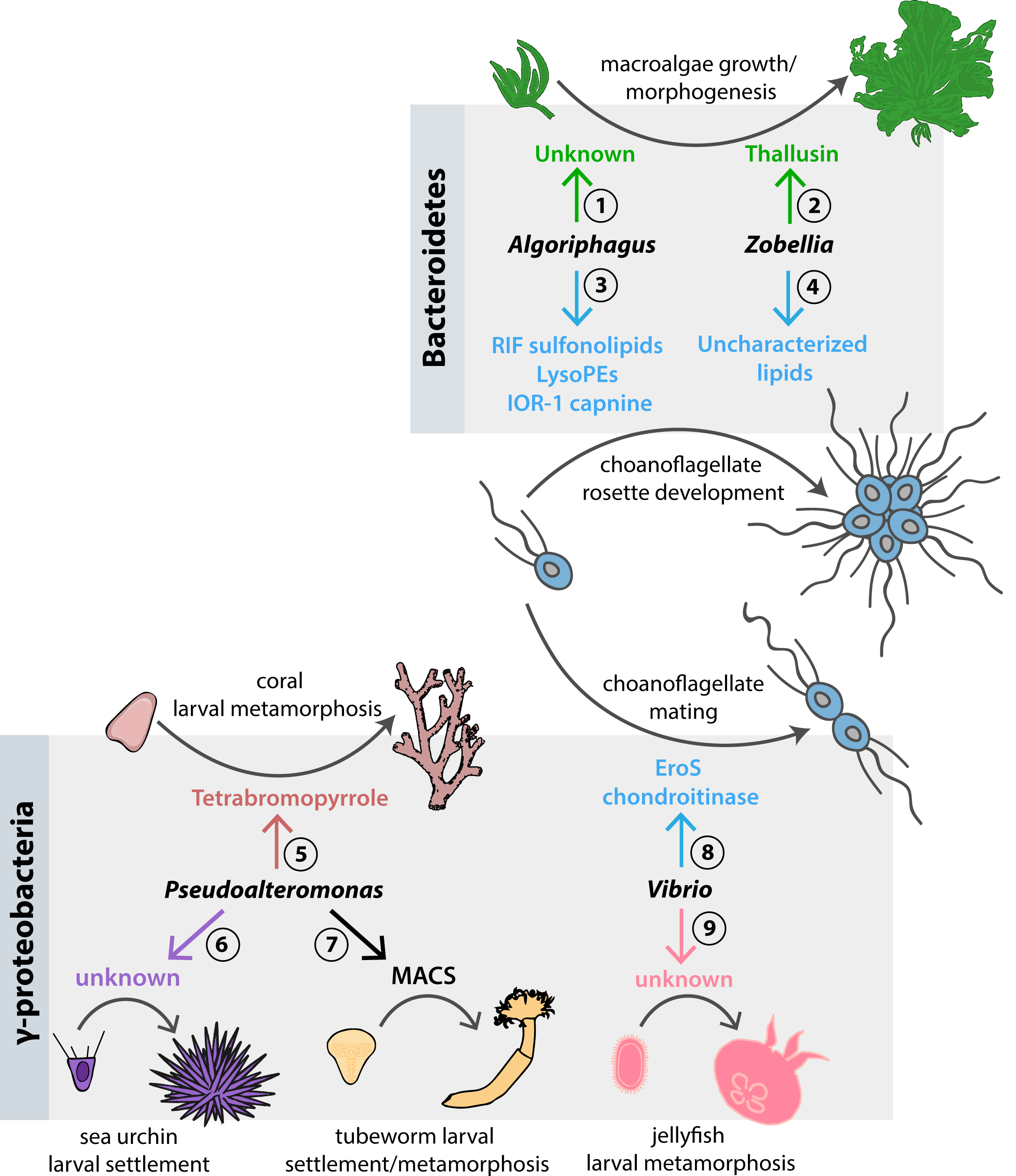
Distinct molecular cues from environmental Bacteroidetes and Gammaproteobacteria regulate developmental transitions in diverse marine eukaryotes. The Bacteroidetes bacteria *Algoriphagus* and *Zobellia uliginosa* regulate morphogenesis in organisms as diverse as algae and choanoflagellates. **(1)** Uncharacterized factors produced by *Algoriphagus* induce morphogenesis in the macroalgae *Ulva mutabilis.* **(2)** *Algoriphagus machipongonensis* lipids [sulfonolipids, lysophosphatidylethanolamines, and a capnine] regulate rosette development in the choanoflagellate *Salpingoeca rosetta*. **(3)** Thallusin, and amino acid derivative produced by *Zobellia uliginosa*, induces morphogenesis in the macroalgae *Monostroma oxyspermum*. **(4)** Uncharacterized molecules from *Zobellia uliginosa* induce rosette development in the choanoflagellate *Salpinogeca rosetta*. Gammaproteobacteria can likewise elicit developmental responses in diverse animals and choanoflagellates. **(5)** Tetrabromopyrrole produced by *Pseudoalteromonas* spp. induces larval metamorphosis in corals *Acropora millepora* and *Acropora willisae*, and larval settlement (attachment and metamorphosis) in the coral *Porites astreoides*. **(6)** Uncharacterized cues from *Pseudoalteromonas* bacteria induce larval settlement in the sea urchin *Heliocidaris erythrogramma*. **(7)** *Pseudoalteromonas luteoviolacea* produces arrays of contractile phage tail-like structures (MACs) that trigger metamorphosis of the tubeworm *Hydroides elegans*. **(8)** A chondroitinase (EroS) secreted by *Vibrio fischeri* induces mating in the choanoflagellate *S. rosetta.* **(9)** Unknown cues secreted by *Vibrio alginolyticus* induce larval metamorphosis in the jellyfish *Cassiopea andromeda*

Because *S. rosetta* and *Algoriphagus* could be cultured independently or together, and because rosette development was quantifiable (i.e. % of cells in rosettes), a straightforward rosette development bioassay could be used to investigate the molecular basis of *Algoriphagus* rosette-inducing activity. Activity-guided fractionation led to the isolation of RIF-1 (<u>R</u>osette-Inducing <u>F</u>actor-1), a novel sulfonolipid signaling molecule that induced rosette development in *S rosetta* [11]. However, only a small fraction of *S. rosetta* cells formed rosettes in response to RIF-1, far fewer than that induced by live *Algoriphagus,* leading to the hypothesis that additional *Algoriphagus* molecules influence *S. rosetta* rosette development [12].

Further work revealed that *Algoriphagus* produces additional lipid activators, synergistic enhancers, and inhibitors that regulate rosette development [12,13] (Fig. 1a). While the RIFs (RIF-1 and a second sulfonolipid, RIF-2) were sufficient to induce low levels of rosette development, an additional class of lipid synergists, the lysophosphatidylethanolamines (LPEs), were required for robust rosette induction.

Together, the RIFs and LPEs recapitulated the full rosette inducing activity of live *Algoriphagus*. The importance of the LPEs had initially been obscured by the fact that they did not exhibit any bioactivity on their own; only by testing bacterial lipid fractions in combination with the RIFs did it become clear that these synergistic lipids helped to fully potentiate the induction of rosette development. Testing bacterial fractions in combination also revealed that *Algoriphagus* produces a molecule that competes with and inhibits RIF-induced rosette development. The molecule, a capnine called IOR-1 (Inhibitor <u>o</u>f <u>R</u>osettes-1), antagonizes the RIFs, but its inhibitory activity can be bypassed in the presence of LPEs, providing a possible explanation for why IOR-1 does not normally prevent *Algoriphagus* rosette induction.

Why might *S. rosetta* rely on a network of bacterial cues before committing to rosette development? We hypothesize that requiring multiple bacterial cues may ensure that rosette development is not initiated in response to the wrong bacteria, or under unfavorable environmental conditions. This integrated response may be especially important in aquatic environments, where bacterial composition and nutrient availability are constantly changing.

### A bacterial chondroitinase triggers mating in S. rosetta

In addition to rosette development, *S. rosetta* can transition from asexual proliferation to sexual reproduction, wherein solitary haploid cells fuse to produce a diploid cell that will then undergo meiosis [14]. Despite harboring a complete meiotic genetic toolkit [15,16], the *S. rosetta* sexual cycle was rarely observed in laboratory cultures. Only under starvation conditions would a small fraction of the *S. rosetta* population mate [17]. A serendipitous observation revealed that specific environmental bacteria, missing from most laboratory cultures, were capable of triggering a robust, population-wide switch to sexual reproduction [18].

This discovery stemmed from the observation that *Vibrio fischeri,* an abundant marine bacterium, induced the formation of large motile aggregates or “swarms” composed of many solitary *S. rosetta* cells. Swarming had not been previously described in *S. rosetta*, and further examination revealed that during swarming, haploid *S. rosetta* cells frequently paired off and underwent cell and nuclear fusion. Genetic experiments confirmed that the diploid products of cell and nuclear fusion later generated meiotic progeny, demonstrating that *Vibrio* bacteria induce the full sexual cycle in *S. rosetta* (Figure 1b).

Because swarming was always observed prior to mating, *S. rosetta* swarming provided a robust bioassay for identifying the molecular basis of the *Vibrio* “aphrodisiac” activity. Activity-guided fractionation led to the isolation of a protein, named EroS (<u>E</u>xtracellular <u>R</u>egulator <u>o</u>f <u>S</u>ex) that fully recapitulated the activity of *Vibrio* bacteria.

Biochemical assays revealed that EroS belongs to a class of bacterial polysaccharide-degrading enzymes called chondroitinases, and that the chondroitin-degrading activity of EroS is sufficient to induce mating in *S. rosetta*. Finally, the *S. rosetta* target of EroS was identified as the sulfated polysaccharide chondroitin sulfate, a component of the extracellular matrix previously thought to be restricted to the animal lineage. As the first example of an environmental bacterium regulating eukaryotic sexual reproduction, the interaction between *Vibrio* and *S. rosetta* raises the possibility that mating in other aquatic eukaryotes may be influenced by environmental bacteria as well.

### Bacteria as master regulators of S. rosetta life history in the marine environment

Bacteria are required for rosette development and mating under laboratory conditions – but can bacteria plausibly regulate *S. rosetta* development in nature? Despite their underlying molecular differences, the cues that induce rosette development and mating are bioactive at environmentally relevant concentrations. The purified *Algoriphagus* RIFs and LPEs display activity at high nanomolar to low micromolar concentrations in the laboratory; yet, the hydrophobicity of these molecules makes it unlikely that *S. rosetta* encounters them as isolated lipids in the environment. As constituents of the *Algoriphagus* outer membrane, it is more likely that RIFs and LPEs are released into the environment within outer membrane vesicles (OMVs), spherical packages of periplasmic content constitutively produced by Gram negative bacteria [19,20]. Indeed, *Algoriphagus* OMVs elicit robust rosette development [12], and retain their bioactivity under a wide range of conditions. Moreover, diverse bacteria belonging to several marine Bacteroidetes and Actinobacteria genera induce rosette development in *S. rosetta* ([11]; unpublished data]), raising the likelihood that *S. rosetta* might encounter rosette-inducing bacteria in multiple environments

In contrast with the lipid regulators of rosette development, the mating-inducing chondroitin lyase, EroS, is a soluble protein constitutively secreted by *Vibrio* bacteria. Not only does EroS trigger mating at picomolar concentrations, but *S. rosetta* swarms in response to as few as 400 *V. fischeri* cells/mL—a density similar to that of *V. fischeri* in oligotrophic oceans [21]. In addition to *V. fischeri,* several other species of *Vibrio* bacteria induce mating in *S. rosetta*, as do commercial chondroitin lyases isolated from *Flavobacterium heparinum* and *Proteus vulgaris* [18], suggesting that encounters between *S. rosetta* and molecules produced by mating-inducing bacteria might be common occurrences in the ocean.

Thus, it is reasonable to infer that *S. rosetta* comes across both rosette development and mating inducing bacteria in nature. In addition, because *S. rosetta* can respond to cues from diverse bacteria, it seems plausible that other life history transitions in choanoflagellates, such as settlement (the attachment of a planktonic cell to a substrate; [6,10]), are regulated by environmental bacteria as well.

## The widespread influences of environmental bacteria

Choanoflagellates are not the only eukaryotes taking life advice from environmental bacteria. Environmental bacteria also regulate developmental transitions in diverse marine algae and animals, and simple model systems are beginning to uncover the bacterial cues that influence eukaryotic morphogenesis.

### Algal morphogenesis

It has long been known that microbial communities associated with the surfaces of marine macroalgae are essential for their growth and morphogenesis [22]. Yet, the bacterial species responsible for stimulating algal development remained elusive for many years, due to the complex and seasonally-shifting composition of algal-associated bacterial communities [23]. A key advance in studying bacterial-algal interactions was the development of axenic culturing techniques (including for the seaweed *Monostroma oxyspermum*), which provided a platform for testing individual bacterial species for morphogenesis-inducing activity [24]. Although diverse bacteria that influence algal growth and morphogenesis have now been identified (Table 1), only one morphogenetic factor, Thallusin, has been isolated by activity-guided fractionation and characterized to date [25–27]. Thallusin is an amino acid derivative produced by the *Monostroma*-associated bacterium, *Zobellia uliginosa*, that is sufficient to induce thallus development in *M. oxyspermum* and partially promote thallus development in *Ulva* species. With a clear bioassay available, why haven’t more algal morphogenetic factors been isolated? One hypothesis is that the bacterial cues regulating algal growth and morphogenesis are produced at very low levels. This was certainly true for Thallusin, although it was ultimately possible to isolate the molecule because of its potency and stability [26].

**Table 1.**
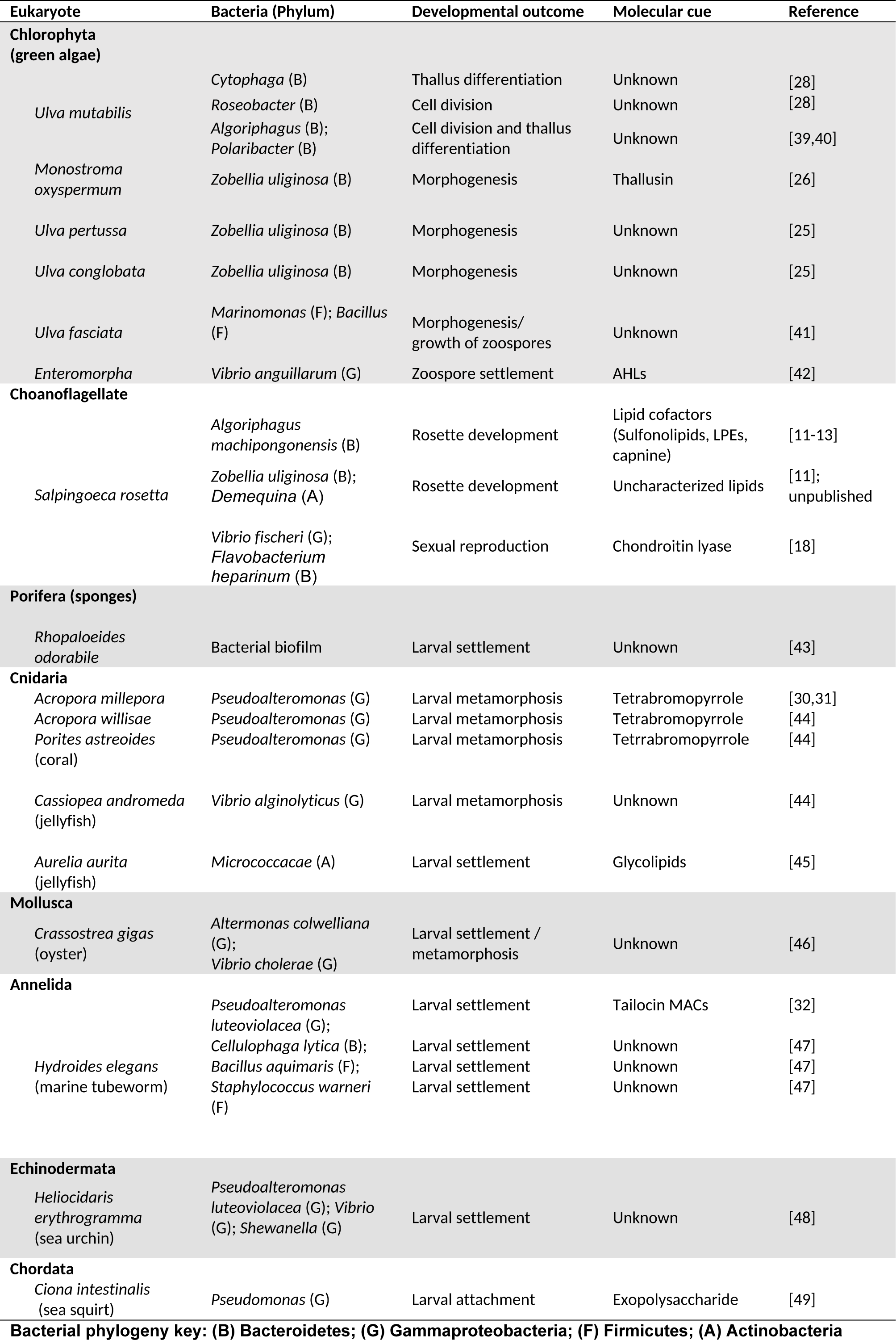
Environmental bacteria regulate development in marine eukaryotes

| Eukaryote | Bacteria (Phylum) | Developmental outcome | Molecular cue | Reference |
| --- | --- | --- | --- | --- |
| **Chlorophyta (green algae)** |  |  |  |  |
| *Ulva mutabilis* | *Cytophaga* (B) | Thallus differentiation | Unknown | [28] |
|  | *Roseobacter* (B) | Cell division | Unknown | [28] |
|  | *Algoriphagus* (B);<br>*Polaribacter* (B) | Cell division and thallus differentiation | Unknown | [39,40] |
| *Monostroma oxyspermum* | *Zobellia uliginosa* (B) | Morphogenesis | Thallusin | [26] |
| *Ulva pertusa* | *Zobellia uliginosa* (B) | Morphogenesis | Unknown | [25] |
| *Ulva conglobata* | *Zobellia uliginosa* (B) | Morphogenesis | Unknown | [25] |
| *Ulva fasciata* | *Marinomonas* (F); *Bacillus* (F) | Morphogenesis/<br>growth of zoospores | Unknown | [41] |
| *Enteromorpha* | *Vibrio anguillarum* (G) | Zoospore settlement | AHLs | [42] |
| **Choanoflagellate** |  |  |  |  |
| *Salpingoeca rosetta* | *Algoriphagus machipongonensis* (B) | Rosette development | Lipid cofactors (Sulfonolipids, LPEs, capnine) | [11-13] |
|  | *Zobellia uliginosa* (B); *Demequina* (A) | Rosette development | Uncharacterized lipids | [11]; unpublished |
|  | *Vibrio fischeri* (G); *Flavobacterium heparinum* (B) | Sexual reproduction | Chondroitin lyase | [18] |
| **Porifera (sponges)** |  |  |  |  |
| *Rhopaloeides odorabile* | Bacterial biofilm | Larval settlement | Unknown | [43] |
| **Cnidaria** |  |  |  |  |
| *Acropora millepora* | *Pseudoalteromonas* (G) | Larval metamorphosis | Tetrabromopyrrole | [30,31] |
| *Acropora willisae* | *Pseudoalteromonas* (G) | Larval metamorphosis | Tetrabromopyrrole | [44] |
| *Porites astreoides* (coral) | *Pseudoalteromonas* (G) | Larval metamorphosis | Tetrabromopyrrole | [44] |
| *Cassiopea andromeda* (jellyfish) | *Vibrio alginolyticus* (G) | Larval metamorphosis | Unknown | [44] |
| *Aurelia aurita* (jellyfish) | *Micrococcaceae* (A) | Larval settlement | Glycolipids | [45] |
| **Mollusca** |  |  |  |  |
| *Crassostrea gigas* (oyster) | *Altermonas colwelliana* (G);<br>*Vibrio cholerae* (G) | Larval settlement / metamorphosis | Unknown | [46] |
| **Annelida** |  |  |  |  |
| *Hydroides elegans* (marine tubeworm) | *Pseudoalteromonas luteoviolacea* (G); | Larval settlement | Tailocin MACs | [32] |
|  | *Cellulophaga lytica* (B); | Larval settlement | Unknown | [47] |
|  | *Bacillus aquimaris* (F); | Larval settlement | Unknown | [47] |
|  | *Staphylococcus warneri* (F) | Larval settlement | Unknown | [47] |
| **Echinodermata** |  |  |  |  |
| *Heliocidaris erythrogramma*<br>(sea urchin) | *Pseudoalteromonas luteoviolacea* (G); *Vibrio* (G); *Shewanella* (G) | Larval settlement | Unknown | [48] |
| **Chordata** |  |  |  |  |
| *Ciona intestinalis*<br>(sea squirt) | *Pseudomonas* (G) | Larval attachment | Exopolysaccharide | [49] |

**Bacterial phylogeny key: (B) Bacteroidetes; (G) Gammaproteobacteria; (F) Firmicutes; (A) Actinobacteria**

Alternatively, induction may require multiple bacterial molecules. On their own, bacteria belonging to *Cytophaga* and *Roseobacter* genera induce incomplete *Ulva mutabilis* development, promoting either cell division or thallus differentiation, respectively [28].

However, the combined activities of these bacteria fully restore normal morphogenesis, raising the possibility that the synergistic interactions observed at the organismal level are required at the molecular level as well.

### Larval settlement

Many benthic marine invertebrates have complex life histories that include stages of larval settlement and metamorphosis, key developmental steps that are crucial for adult success. Bacterial biofilms provide cues that trigger larval settlement and metamorphosis in diverse marine invertebrates, including sponges, cnidarians, molluscs, annelids, echinoderms, and urochordates (Table 1). While the majority of these interactions remain poorly understood, the increased tractability of invertebrate systems (e.g. the tubeworm *Hydroides elegans* [29] and the coral *Acropora millepora* [30]) has facilitated identification of the bacterial cues that regulate larval settlement and metamorphosis. Biofilm-forming bacteria from the genus *Pseudoalteromonas* influence development in several animals, including corals and tubeworms. Interestingly, the cues produced by environmental *Pseudoalteromonas* that trigger metamorphosis in *A. millepora* and *H. elegans* are distinct; while *A. millepora* metamorphosis is induced by the small molecule tetrabromopyrrole [31], *H. elegans* metamorphosis is regulated by arrays of contractile phage-tail like structures called MACs (<u>M</u>etamorphosis <u>A</u>ssociated <u>C</u>ontractile <u>S</u>tructures) [32]. Although the structures of tetrabromopyrrole and MACs suggest that the mechanisms by which these molecules trigger metamorphosis are likely very different, both of these bacterial molecules may provide chemical evidence of a suitable surface for colonization. Because surfaces in the ocean are often limiting, cues from bacteria might indicate to animals that they have found an appropriate environment for settling down.

## Bacterial cues are proxies for environmental conditions

As more bacterial cues are isolated, it is becoming clear that interactions between eukaryotes and their environmental bacteria exhibit remarkable molecular specificity. Even slight modifications to the structures of bacterial cues can completely eliminate inducing activity, as is the case with the choanoflagellate rosette-inducing molecules and the algal morphogenetic factor Thallusin [13,27,33]. Nonetheless, multiple environmental bacteria can elicit the same eukaryotic developmental responses, and in each case the molecular cues seem to be distinct (this has been demonstrated for *S. rosetta* rosette development, *Monostroma* morphogenesis, and *H. elegans* larval settlement; Table 1). Because marine microbial communities are highly dynamic, it may be beneficial for eukaryotes to interpret developmental cues from diverse bacteria. The molecular stringency we observe likely allows eukaryotes to be responsive to many different environmental bacteria, whilst maintaining tight regulation over important developmental decisions.

Interestingly, we find that select bacterial genera (including *Flavobacteriia, Pseudoalteromonas,* and *Vibrio*) have a high level of influence on the development of diverse marine eukaryotes. It is possible that many eukaryotes may rely on cues from Bacteroidetes and Gammaproteobacteria because these bacteria flourish in nutrient and carbon rich environments, and can thus serve as proxies for favorable environmental conditions. Indeed, many of the bacteria that regulate eukaryotic development are well equipped for rapidly responding to increasing nutrient availability, boasting an assortment of extracellular enzymes that break down polysaccharides, lipids, and proteins [34]. For example, the polysaccharide degrading abilities of Bacteroidetes and Gammaproteobacteria allow these bacteria to utilize algal-derived polysaccharides and promptly proliferate when phytoplankton bloom (often as a result of increased mineral levels) [35]. Even more impressive is the ability of some inducing-bacteria (notably *Vibrio* spp.) to pursue nutrient dense microenvironments through chemotaxis [36]. Thus, diverse eukaryotes may have converged on certain bacteria as indicators of nutrient-rich environments.

Finally, it is enticing to consider how environmental bacteria might benefit from these interactions. Many of the bacteria that induce eukaryotic developmental transitions also frequently associate with eukaryotes, for example by accumulating on surfaces of macroalgae and invertebrates. Because these bacteria produce exoenzymes that help them utilize plant and animal-derived molecules for nutrition [34], it is possible that inducing eukaryotic development allows specific bacteria to rapidly colonize valuable “real estate.”

## Conclusion

Although the influences of environmental bacteria on the development of marine eukaryotes has been observed for decades, we are just beginning to gain a molecular understanding of these interactions. With the help of simple model systems and straightforward bioassays, it is becoming clear that environmental bacteria produce structurally diverse cues that govern eukaryotic development with a high degree of molecular specificity. Nonetheless, even simple bacterial-eukaryotic interactions can be challenging to characterize when we rely solely on activity-guided fractionation. Furthermore, most interactions with bacteria are not simple, and may rely upon multiple molecular cues produced by one or more bacterial species. To understand these complex interactions, future studies should likely combine genetic and activity-guided approaches in the study of environmentally-relevant pairs or communities of eukaryotes and bacteria.

Importantly, many of the environmental bacteria that regulate eukaryotic development in marine ecosystems interact with eukaryotes in other pathogenic and commensal contexts. For example, the bacterium *Vibrio fischeri* forms a symbiosis with and triggers morphogenesis in the Hawaiian bobtail squid (*Euprymna scolopes*), while other members of the Vibrionaceae are common animal enteric commensals, pathogens, and mutualists [37]. In addition, Bacteroidetes bacteria closely related to

*Algoriphagus* are abundant mammalian gut commensal bacteria that are important for proper intestinal development and homeostasis [38]. Therefore, understanding how environmental bacteria shape the development of marine eukaryotes may also provide insight into broadly applicable mechanisms of bacterial-eukaryotic interactions.

## Acknowledgements

We thank D. Booth, E. Ireland, B. Larson, T. Linden providing feedback on the review prior to submission. Research in the King laboratory is supported by the Howard Hughes Medical Institute, the National Institutes of Health (R01GM099533), and the Gordon and Betty Moore Foundation.

